# Orthogonal but linked neural codes for value

**DOI:** 10.1101/2021.07.27.453966

**Authors:** David J-N. Maisson, Justin M. Fine, Seng Bum Michael Yoo, Tyler V. Cash-Padgett, Maya Zhe Wang, Brianna J. Sleezer, Jan Zimmermann, Benjamin Y. Hayden

**Affiliations:** Department of Neuroscience, Center for Magnetic Resonance Research, Center for Neuroengineering, Department of Biomedical Engineering University of Minnesota, Minneapolis MN 55455

**Keywords:** Expectancy, value, reward, abstract value, neuroeconomics

## Abstract

Our ability to effectively choose between dissimilar options implies that information regarding the options’ values must be available, either explicitly or implicitly, in the brain. Explicit realizations of value involve single neurons whose responses depend on value and not on the specific features that determine it. Implicit realizations, by contrast, come from the coordinated action of neurons that encode specific features. One signature of implicit value coding is that population responses to offers with the same value but different features should occupy semi- or fully orthogonal neural subspaces that are nonetheless linked. Here, we examined responses of neurons in six core value-coding areas in a choice task with risky and safe options. Using stricter criteria than some past studies have used, we find, surprisingly, no evidence for abstract value neurons (i.e., neurons with the response to equally valued risky and safe options) in any of these regions. Moreover, population codes for value resided in orthogonal subspaces; these subspaces were linked through a linear transform of each of their constituent subspaces. These results suggest that in all six regions, populations of neurons embed value implicitly in a distributed population.

## INTRODUCTION

Humans and other animals can readily compare options that differ qualitatively - even if those options are dissimilar. Our fluency with choices between distinct options has led many to conclude that there exists a single value scale (Tremblay & Schultz, 1999; Elliott et al., 2000; Levy & Glimcher, 2011; Padoa-Schioppa, 2011; McNamee et al., 2013; O’Doherty, 2014). The question of how this value scale is realized in the brain has been central to the field of neuroeconomics (Rangel et al., 2008; Kable & Glimcher, 2009; Rushworth et al., 2011; Wunderlich et al., 2012; Padoa-Schioppa & Conen, 2017). One view, which we call the explicit view, holds that value is encoded in the firing rate responses of single neurons; such neurons can be said to be tuned for value. Previous studies have indeed shown evidence consistent with this hypothesis, namely that there are neurons, located within core reward regions of the brain, that have firing rates that covary with the values of offers or choices regardless of the features that determine those options’ values (Platt & Glimcher, 1999; Tremblay & Schultz, 1999; Kennerley & Wallis, 2009; Cai et al., 2011; Padoa-Schioppa, 2011; Kim et al., 2012; Farashahi et al., 2019; Padoa-Schioppa & Conen, 2017; Levy & Glimcher, 2011, 2012; Bartra et al., 2013). These hypothesized value neurons would necessarily respond identically to two equally preferred options no matter how different their features. Indeed, this pattern of firing would be how such options are compared.

A second view, which we call the implicit view, holds that value is instead coded implicitly – that is, not in the responses of single neurons, but as an emergent feature of populations of neurons, none of which is selective to value *per se*. Such a value code would have to be compositional - that is, it would depend critically on the coordinated activity of multiple individual neurons whose responses are not themselves necessarily value-specific. In such a framework, neural representations of value would be flexibly composed from the responses of neurons that encode mixtures of the features underlying value (Rigotti et al., 2013; Fusi et al., 2016; Bernardi et al., 2020). Such neurons would presumably encode the features of the anticipated outcome; that is, they would encode outcome expectancies (Schoenbaum et al., 1998, 2003, 2009, 2011; Zhou et al., 2021). The activation of a population of such neurons could, when appropriately combined, result in an implicit (or emergent) encoding of abstract value. Such coding could permit comparison of dissimilar but equally valued options even without abstract value neurons.

While the evidence may seem to solidly favor the existence of pure value neurons, it turns out to be quite difficult to eliminate confounding factors, so the issue is still hotly debated (Maunsell, 2004; Kable & Glimcher, 2009; Wallis & Rich, 2011; O’Doherty, 2014; Stalnaker et al., 2015; Hayden & Niv, 2021; Zhou et al., 2021). We propose two hypotheses that differentiate implicit from explicit single neuron coding of value. The ***first hypothesis*** is that similarly valued options with qualitatively different features will not match, but instead have distinct neural codes. The existence of such codes would argue against the existence of an abstract value code, which requires subjectively equal values to be encoded the same way. Indeed, they would have to be in the same subspace, whereas implicit coding implies that neural responses to qualitatively distinct offers will occupy distinct subspaces. Consequently, the ensemble response will occupy a different part of the larger response manifold (Tang et al., 2020; Russo et al., 2020; Cohen et al., 2021; Ebitz & Hayden, 2021; Panichello & Buschman, 2021). The first hypothesis directly implies the ***second hypothesis***, namely, that if different options occupy separate subspaces due to their different features, those subspaces must somehow be linked by some lawful transformation to support a value comparison process for driving the ultimate decision (Elsayed et al., 2016).

The idea of separate-but-linked subspaces was originally proposed in the motor system; pre- and peri-movement activity are in separate subspaces (Kaufman et al., 2014), but those subspaces are lawfully linked (Elsayed et al., 2016). We have also shown that these relationships apply outside the motor system, to core reward regions, albeit in a completely different context: to temporally separated evaluation and comparison periods in a choice task (Yoo & Hayden, 2020). Here, we hypothesize that transformations of separate value subspaces into a common subspace provide a mechanism for comparison of dissimilar options.

For our dissimilar goods, we focused on a well-studied paradigm, the choice between risky and safe options (McCoy & Platt, 2005; Platt & Huettel, 2008; Kim et al., 2012; So & Stuphorn, 2016). It is well established in the psychological literature that such options differ, not simply in their variance, but in the psychological processes they evoke (Weaver, 1982; Lopes, 1987; Loewenstein et al., 2001; Barseghyan et al., 2013). Nonetheless, humans and monkeys can adroitly compare risky and safe offers to each other in an economically meaningful way (Heilbronner & Hayden, 2013; Petrillo et al., 2015; Heilbronner, 2017). We used a risky choice task to investigate neuronal responses in six core value regions, the ventromedial prefrontal cortex (vmPFC, area 14), orbitofrontal cortex (OFC, area 13), rostral OFC (rOFC, area 11), pregenual anterior cingulate cortex (pgACC, area 32), posterior cingulate cortex (PCC, area 29), and ventral striatum (VS). In all six areas, we found that responses evoked by equally valued risky and safe options differed and, indeed, occupied distinct subspaces. The population response subspaces were linked in such a way that their separate subspaces could be aligned through a rotational transformation.

## RESULTS

### Behavior

On each trial of the *risky choice task* (Strait et al., 2014) (see **Methods**), a macaque (*Macaca mulatta*) subject chose between two offers that varied in magnitude and probability (**Figure 1A**). Safe offers (12.5% of offers) provided a small volume of juice (125 *μ*L) with 100% certainty. Risky offers provided either a medium (165 *μ*L, 43.75% of offers) or large (240 *μ*L, 43.75% of offers) volume of juice with a defined probability (0-100%, 1% increments). The offer types for the two offers were selected by a computer at random and independently on each trial. Our dataset consists of responses from six subjects in 315 recording sessions and consists of 211,884 trials (average 672.6 trials per session). Target regions for neural recordings are illustrated in **Figure 1B** and anatomical boundaries are provided in the **Methods**.

**Figure 1.**
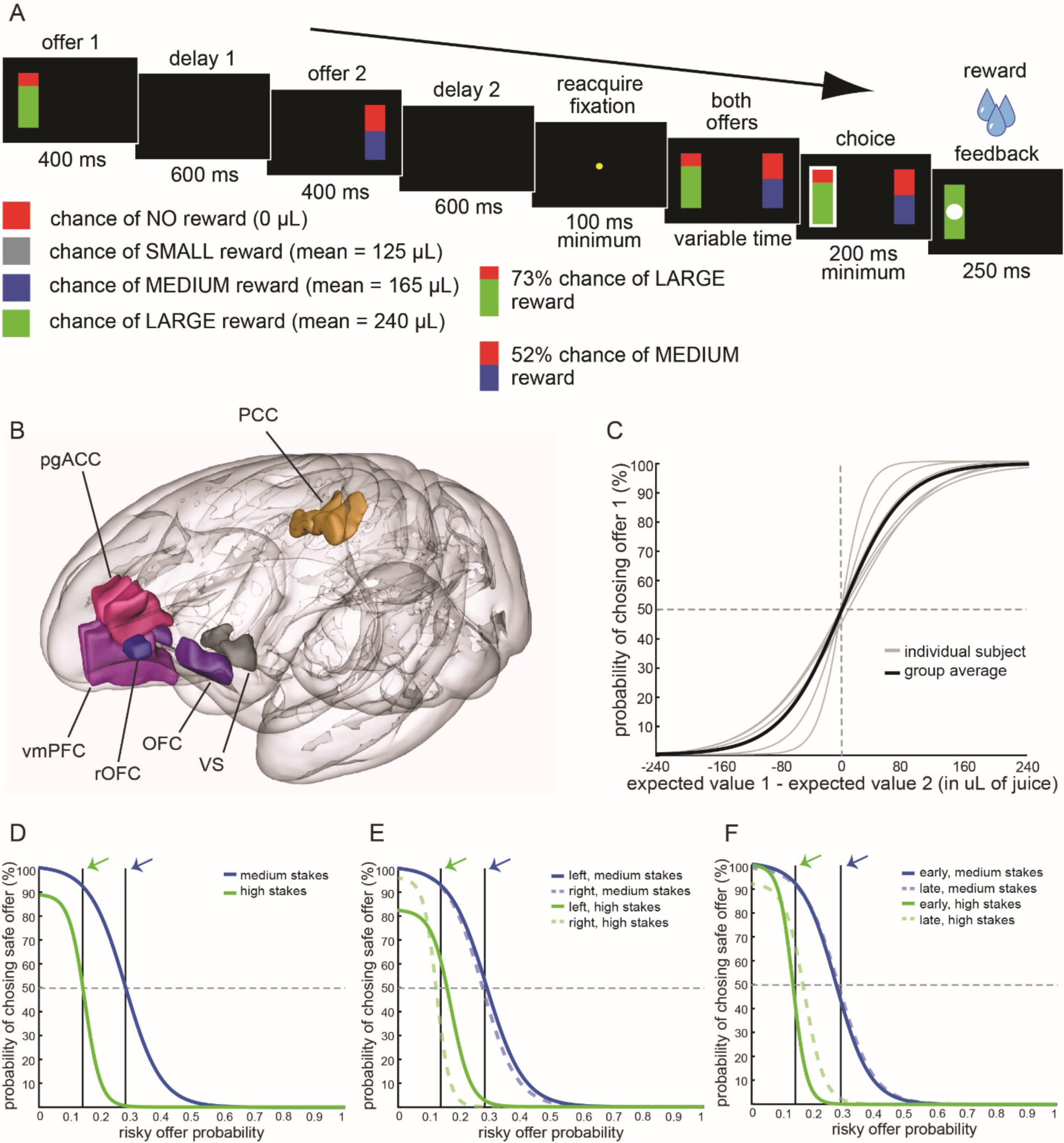
Task, targeted structures, behavior, and calculation of equivalent risky and safe values. **(A)** Structure of our Risky Choice Task (Strait et al., 2014). Each trial begins with a 400 ms presentation of the first offer followed by a 600 ms blank period. Following a 400 ms presentation of the second offer and another 600 ms blank period, a fixation spot appears and, on fixation, both offers appear and the subject selects one by saccade. For each offer, the magnitude of the associated reward (stakes) is indicated by the bottom color (green, high or blue, medium) of the stimulus. The probability of being rewarded is indicated by the size of the green/blue segment. **(B)** Anatomical positions of our brain regions of interest: rOFC (blue), OFC (purple), vmPFC (purple-pink), pgACC (pink), PCC (gold), and VS (grey). **(C)** Likelihood of choosing the first offer as a function of its value relative to the second (specifically, for signed value difference). Sigmoid fits of raw binary data shown (see **Methods**). Gray lines: individual subjects; black line: group average. In this and subsequent panels, a horizontal, dashed line indicates the indifference point (the point at which choices are 50/50). **(D)** Likelihood of choosing a safe option as a function of the probability of the risky option for medium (blue) and high (green) stakes offers. All data were analyzed on a subject-by-subject basis, so only data for one example subject (subject B) are shown. Other subjects showed similar patterns. Vertical black lines indicate the probability used as the SV-equivalence point for the subject (the arrow points to the indifference point for medium (blue) and high magnitude (green) risky offers). **(E)** Same as D, except data are separated for left and right offers. Side of presentation does not affect choice much. **(F)** Same as D, except data are separated by trials that were in the first (early) or second (late) half of a session.

Subjects consistently performed at a high level, were modestly risk-seeking, and did not differ from each other qualitatively (**Figure 1C**). Details of species-typical behavior in this task are given elsewhere (in greatest detail in Hayden et al., 2010; Farashahi et al., 2018, 2019). Results of these analyses are not repeated here, except to confirm, as we have previously shown, that subjects’ behavior is quite stable and consistent within subject, both across and within sessions, and across subjects and sessions **(Figure 1D-F**). Indeed, all subjects showed the same behavioral patterns we have observed using this task in past studies.

All analyses in this paper make use of subjective values instead of expected values. To identify the relative values of safe offers, we computed the risky-safe indifference point (Hayden et al., 2010). To do this, we calculated, separately for each subject and separately for medium and large stakes offers, the likelihood that the subject would choose the safe offer as a function of the probability associated with the risky offer. We then calculated indifference using a standard approach in which we fit the resulting data with a sigmoid curve and calculated the point at which the best-fitting curve crossed the indifference line (see **Methods**). As in our past studies, all subjects were risk-seeking (Heilbronner & Hayden, 2013; Heilbronner, 2017). The cross-subject average indifference point for medium-stakes risky offers corresponded to an offer probability of 0.33 ± 0.05 (standard deviation). A risk-neutral subject would have had an indifference point at 0.76. The average indifference point for high-stakes risky offers corresponded to an offer probability of 0.11 ± 0.04. A risk-neutral subject would have had an indifference point of 0.52. As we have observed many times, preferences were highly consistent across many contexts. For example, indifference points are similar for the first and second offers (offer 1: medium: 0.34; high: 0.11; offer 2: medium: 0.22; high = 0.11), for offers made early and late in the session (early: medium: 0.29; high: 0.09; late: medium: 0.31; high: 0.13), and when risky offers appear on the left or right (left: medium: 0.27, high: 0.11; right: medium: 0.36, high: 0.12). Data for an example subject are shown in **Figure 1D-F**.

Because our central research goal is to ascertain the influence of offer type on firing, we wanted to ensure that our effects were not due to attention or arousal. We therefore examined the relationship between two proxies for attention, looking time and pupil size in four of our subjects (subjects B, H, P, and S; these are the subject for which we had pupil size recorded). We found no detectable relationship. Specifically, during the offer 1 epoch, subjects fixated the first offer for 198.5 ms if it was risky and for 199.1 ms if it was safe. These two are not different (p=0.85 for the group, p-values were the same or higher for all four subjects). During the offer 2 epoch, subjects fixated the second offer for 222.4 ms if it was risky and for 220.9 ms if it was safe. During the choice epoch, we calculated total looking time and, again, found no differences (302.7 ms for chose risky and 302.5 for chose safe, p = 0.093). Nor were any of the individual subjects statistically significant for any of these measures (p > 0.05 in all cases). These results do not appear to be dependent on our choice of epoch; we used longer analysis (1 sec) epochs for each of these analyses and found qualitatively matched results.

During the offer 1 epoch, the pupil size did not differ for risky and safe options in any of the four subjects (p=0.10 for subject S and p>0.5 for the other three subjects). Specifically, relative to the baseline value (defined as 0), risky offer-evoked pupil size was -0.92 z-score units (for all subjects averaged); the corresponding value for safe offers was 0.93 z-score units. Likewise, during the offer 2 epoch, relative to the baseline value (again, defined as 0), risky offer-evoked pupil size was -0.81 z-score units (for all subjects averaged); the corresponding value for safe offers was 0.80 z-score units. These are not different for any subject (p=0.21 for subject H, p=0.39 for subject P, p>0.5 for the other two subjects). Finally, during the choice epoch, relative to the baseline value of 0, risky choice-evoked pupil size was -0.79 z-score units (for all subjects averaged); the corresponding value for safe offers was also 0.79 z-score units. These are not different for any subject (p<0.5 in all cases).

### Responses of single neurons to risky and safe offers

We recorded responses of 981 neurons in 6 brain regions while our subjects performed the risky choice task: vmPFC (area 14, 156 neurons), OFC (area 13, 157 neurons), rOFC (area 11, 138 neurons), pgACC (area 32, 255 neurons), PCC (area 29/31, 151 neurons), and VS (nucleus accumbens, 124 neurons). We recorded in two subjects for all areas, although different subjects were used for the different areas (see **Methods**). Detailed analyses of responses to risky offers were reported previously for vmPFC and VS (Strait et al., 2014, 2015). We have never previously examined neural responses to safe offers.

For these and subsequent analyses, we used each individual’s subjective indifference point. We then defined a range of probabilities (± 2.5%, total range of 5.0%) and treated all offers within that range as being subjectively equivalent to the safe value. We subsequently checked for robustness by repeating the following analyses using a larger range (± 5%, total range of 10%) but because we found no qualitative differences, we do not report those results. We also repeated all analyses using within-session estimates of subjective value (rather than within-subject, across-session). Again, because we found no qualitative differences, we do not report those results.

In our task, offers are presented asynchronously. Thus, to compare safe and equally valued risky offers, we compare responses across trials, and within the first offer epoch, and not within a given trial. To give an example, neuron vmPFC.70 (**Figure 2A**) showed differing selectivity for safe and risky offers during the first offer epoch. Neuron pgACC.17 (**Figure 2B**) showed clear and roughly monotonic tunic for probability. This means that a downstream decoder could readily identify the probability (and thus, value) of a risky offer. However, this scale does not extend to safe offers. Because the safe offer had a subjective value equivalent to 0.28, given the explicit view for value encoding, its mean neural response should have been the same as the response to that offer (0.36 spikes/second). Instead, it evoked a mean response of 1.67 spikes/second (these differ, Student’s t-test, t = 4.31, *p* < 0.001). Additional sample cells, from each targeted area, showed positive and negative monotonic tuning for risky values and unrelated responses to safe offers (**Figure 2C-F**).

**Figure 2.**
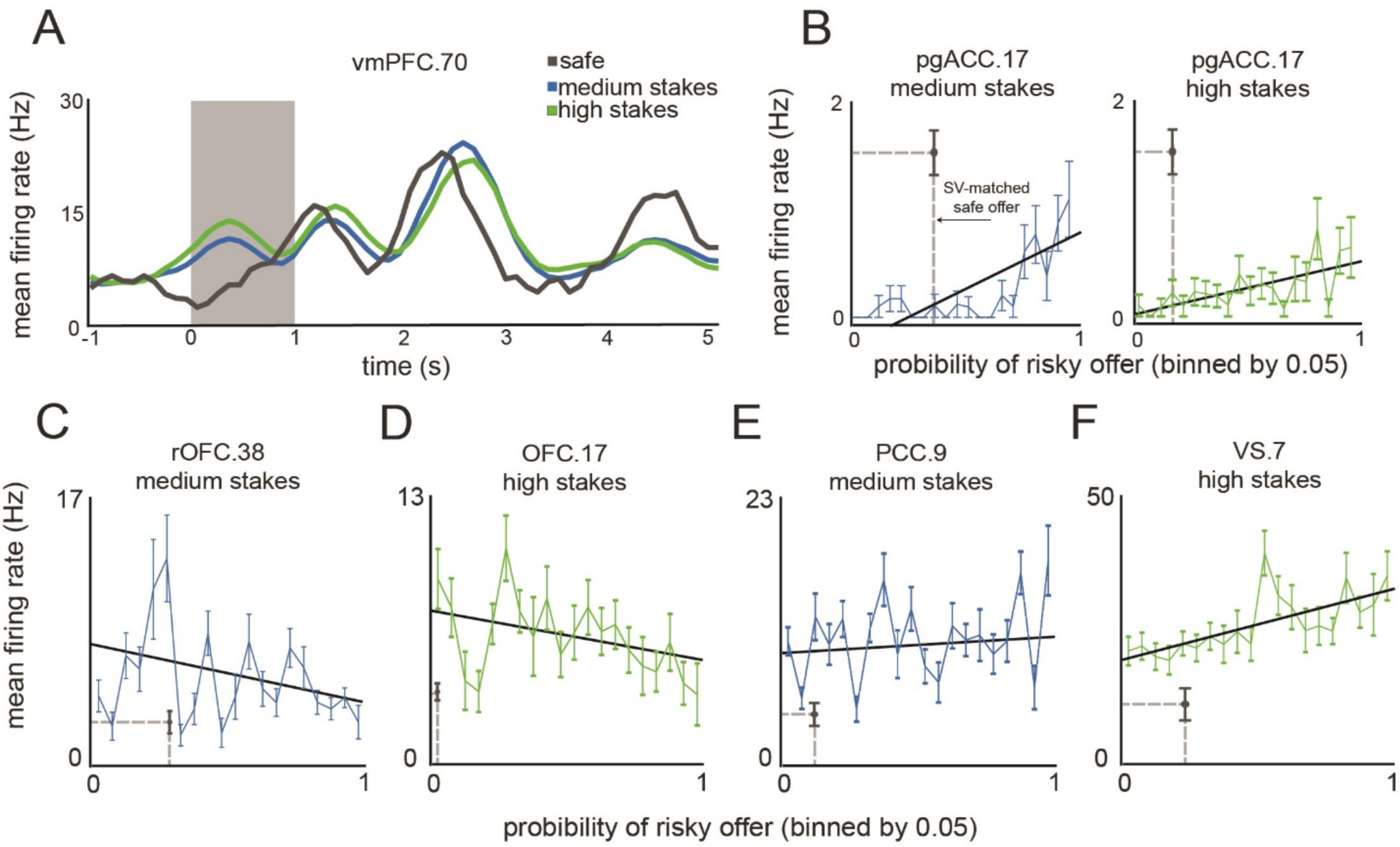
Responses of single neurons. This figure shows the average responses of sample neurons to safe and risky offers of differing values, as well as the average response similarity rates. **(A)** Peristimulus time histogram from mean firing rates of sample neuron vmPFC.70. Each line indicates the average response across offers of a given risk profile (grey: all safe offers; blue: all medium stakes risky offers; green: all large stakes risky offers). The grey shaded box indicates the 1-second period from which the 500-ms epoch 1 analysis window was extracted, where the onset of the first offer is time-locked to zero seconds. **(B)** This is a plot of data collected from a sample neuron pgACC.17, which showed a response to safe offers that was statistically different from the response to equivalent risky offers. Depicted are the average responses to medium stakes (left; blue) and high (right; green) stakes, separated by probability ranges of 0.05. The red point indicates the average response of the given neuron to safe offers (error bars denote the SEM across responses to safe offers). The diagonal black line indicates a fitted regression line, showing positive monotonic tuning. **(C-F)** Same as (B) but demonstrating sample cell responses to an assortment of medium and high stakes offers from across all target areas.

### Responses to safe options are unrelated to responses evoked by equally valued risky options

We wanted to know to what extent the activities of these sample neurons are representative of the population. If the difference between responses to safe vs. matched risky is lower than the difference between responses to random offers, it would indicate that that neuron satisfies a minimal condition for abstract value coding. Alternatively, if the difference is identical, it would be evidence in favor of the idea that this neuron uses distinct or unrelated codes for risky and safe offers. We took advantage of a key feature of our task design: the large range (1-100 by 1% increments) of probabilities tested. Specifically, we compared the difference between safe and equivalent risky options to the difference between safe and randomly selected probability, then repeated this process one thousand times with 1,000 randomly selected probabilities.

In vmPFC, the average medium risky-safe difference was 0.068 (± 0.007 standard deviation) z-score units. (Note that, to give a general cross-neuron measure independent of mean firing, we report results for z-scored data; however, all patterns shown here hold when using raw responses instead). The result in the random condition was nearly identical (0.068 ± 0.002; these were not different, *p* = 0.487, bootstrap test, see Methods). The same pattern is observed in the high stakes risky condition (matched: 0.071; bootstrap: 0.073; not different, *p* = 0.519; **Figure 3A**). The Bonferroni corrected p-values are *p* = 0.698 and *p* = 0.720, respectively. We found the same patterns in the other five structures we examined: none of our six brain areas showed a measurable difference between the safe and random risky offers (summarized in **Figure 3B**).

**Figure 3.**
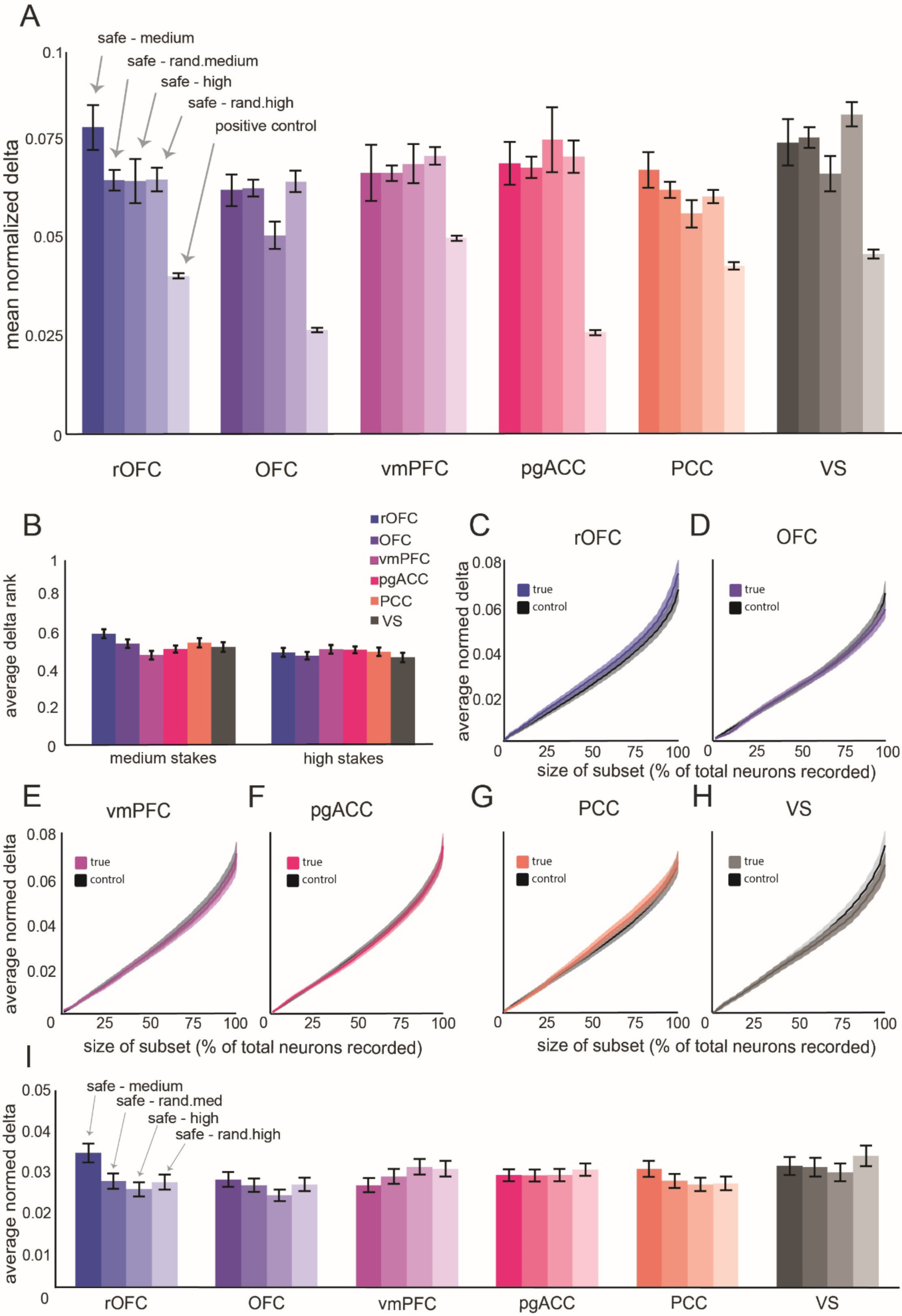
Evoked response difference analysis with best subsets of various sizes. **(A)** The average normalized evoked response difference for each structure. From left to right (most to least opaque), for each structure: evoked response differences in response to safe v. medium magnitude risky offers, safe v. randomly selected medium offer, safe v. high, safe v. random high, and the positive control evoked response differences. **(B)** The average proportion of evoked response differences, across neurons, between random offers that are smaller than the evoked response difference between equivalent safe and risky offers (i.e. the *p-value* from our bootstrap test). **(C-H)** Shown is the change in average evoked response difference (for visualization only: collapsed across both medium and high stakes comparisons) between safe-risky (blue) and safe-random (black). The lines indicate the average normalized evoked response differences (as in panel A), across subsets from a size of 1-100% of recorded neurons. Shaded ribbons denote the standard error across the subset. *Note again that collapsing across medium and high stakes offers is for visualization only. The analysis was performed for safe-medium and safe-high independently. **(I)** For each structure, each bar provides a summary of the average normalized evoked response difference across all subset sizes. Error bars represent the standard error across subset sizes.

The pattern is observed when examining coding of *chosen* options, as opposed to *offered* options. In vmPFC, the average medium risky-safe difference was 0.071 (± 0.006, standard error) z-score units. The result in the random condition was nearly identical (0.076 ± 0.003; these were not different, *p* = 0.311, bootstrap test). The same pattern is observed in the high stakes risky condition (matched: 0.062; bootstrap: 0.067; not different, *p* = 0.462). We found the same patterns in the other five structures we examined (rOFC: medium - 0.081 ± 0.006, *p* = 0.606, high - 0.077 ± 0.006, *p* = 0.607; vmPFC: medium - 0.071 ± 0.006, *p* = 0.523, high - 0.062 ± 0.005, *p* = 0.495; pgACC :medium - 0.081 ± 0.005, *p* = 0.535, high - 0.084 ± 0.006, *p* = 0.584; PCC: medium - 0.088 ± 0.006, *p* = 0.584, high - 0.069 ± 0.004, *p* = 0.565; VS: medium - 0.068 ± 0.005, *p* = 0.544, high - 0.070 ± 0.005, *p* = 0.565).

Our failure to find a difference between the true and random conditions is a null result and could be due to noise in our sample. We therefore developed a *positive control* analysis designed to account for this possibility. We randomly assigned trials from subjective value-matched offers to two sets of equal size, for each neuron. We then computed the risky vs. safe difference for each neuron between these two randomized sets. Because these responses are evoked by stochastically identical stimuli, they must necessarily have zero true difference. Any measured difference between them gives a measure of the difference we would expect to arise by chance. We found that the average medium risky vs. safe difference across random sub-selections, computed from vmPFC responses, was 0.051 z-score units. This value is lower than the safe vs. medium magnitude value in vmPFC (which was 0.068, see above). These values are themselves significantly different (*p* < 0.001, bootstrap test). In other words, we show that we would have detected a pattern consistent with the abstract coding of value if such a code were used; in reality our neurons were more different than could be accounted for by noise. We also found the same pattern for high stakes vs. safe (0.071), which was also significantly greater than the positive control average (0.051, *p* < 0.001). Indeed, we observed the same pattern for medium and high stakes offers in all six structures (*p* < 0.01 in all twelve cases, **Figure 3A**).

We next asked whether there are specialized subpopulations of abstract value-coding neurons. Such a subpopulation, if it exists in an area, must show a lower than chance difference between responses evoked by safe and equivalently valued risky ones (putting aside the possibility of high noise, which we account for below). A putative value-coding population can therefore be identified by taking the subset of neurons with the lowest safe-risky difference. For the full range of possible population sizes (1-100%, by 1% increments), we identified the lowest risky-safe response differences and compared that to the difference observed between safe offer responses and that of a randomly selected risky offer probability; we then repeated this procedure 1000 times. As the size of the tested population increases, naturally so does the average risk-safe difference because we select more neurons with larger response ranges (**Figure 3C-H**). The important comparison is between the safe-risky and safe-random – the explicit view of value encoding predicts that the safe-risky curve will be lower than the safe-random curve. However, in the implicit view, responses to value-matched offers are no more similar than between random offers; the two curves will overlap. For each subsampled population size, we compared the two curves using a Kolmogorov-Smirnov test. We found no significant differences (*p* > 0.19 in all cases; the average normalized evoked response difference across population subsets is summarized in **Figure 3I**).

The same pattern of results holds as well when considering *chosen* value instead of *offered* value (during the choice epoch, see **Methods**). In all six brain regions, we found that there were no significant differences (rOFC: medium - *p* = 0.894, high - *p* = 0.344; OFC: medium - *p* = 0.999, high - *p* = 0.556; vmPFC: medium - *p* = 0.677, high - *p* = 0.677; pgACC: medium - *p* = 1, high - *p* = 0.677; PCC: medium - *p* = 0.961, high - *p* = 0.894; VS: medium - *p* = 0.992, high - *p* = 0.961) between the subpopulation risk-safe differences for safe and matched- risky chosen offer values and those between safe and random-risky chosen offer values.

As with the above analysis, this result relies on a null result. We therefore again applied the logic of a positive control to provide interpretational validity for these results. As above, we reasoned that if value was coded abstractly, then any difference between risk and safe responses is due to noise. And as above, for each neuron, we randomly separated the trials into two sets. We computed the average risk-safe response difference between the two sets. We repeated this random separation 1000 times and computed the average difference across iterations. Then, across neurons, we sorted the resulting differences from least to greatest and identified the subset of neurons (1 - 100%) with the lowest differences. Finally, we performed a Kolmogorov- Smirnov test to compare this positive control distribution to the safe-risky response difference distributions across subset sizes. In OFC, for example, we found that the average difference across positive control subsets was 0.0002 ± 0.00004 (standard error). Both the safe-medium and safe-high risky differences across subsets were significantly different from the positive control differences (*p* < 0.001, Kolmogorov-Smirnov test). The same pattern was true across the remaining 5 areas for both medium and high stakes gambles (*p* < 0.001 in all twelve cases). In other words, the difference between neural response evoked by safe and equally valued risky offers was greater than could be accounted for by noise, indicating that coding schemes are indeed different.

### Offer type (safe vs. risky) is decodable in all six regions

If value is encoded abstractly at the single neuron level; it should not be possible, in principle, to distinguish safe from equally valued risky offers based purely on population neural responses. If, conversely, the brain uses distinct codes for the two categories, then the category (risky vs. safe) should be readily decodable even if the offers are equally valued. We trained a binary support vector machine (SVM) to decode safe from equally valued risky offers based on neural responses (see **Methods**). We then cross-validated the trained model by using it to predict risky vs. safe from responses in a sequestered set. In OFC, for example, we found that the classifier could readily disambiguate safe from equally valued medium-stakes risky offers with 96.9% accuracy, which is significantly greater than the 49.9% decoding accuracy in the shuffle data (t = 730.1, *p* < 0.001; high magnitude: 96.6%, t = 780.5, *p* < 0.001). We also observed clear decodability in the other five structures (*p* < 0.001 in all cases; **Figure 4A**).

**Figure 4.**
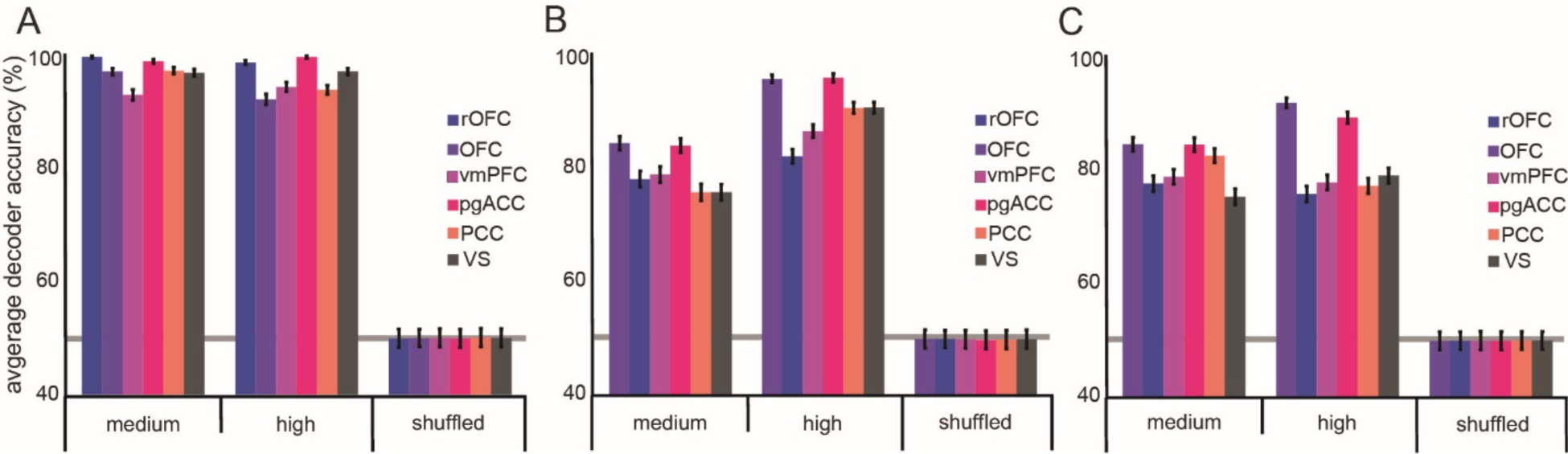
Safe and equivalently valued risky offers are readily decoded. **(A)** Decodability for safe and risky offers is high in all areas for both medium and high stakes gambles. Shuffled data refers to decodability of randomly assigned safe/risky labels to neural responses that are completely shuffled across trials and cells. Error bars indicate the standard error across cross-validations. **(B)** same as A, except that the only neurons used in the decoder were those which showed no significant differences in neural responses to safe and equivalent risky offers, and which have differences in responses within the lowest half of the set. **(C)** same as B, except that the only neurons used are those with differences in responses within the lowest quarter of the set.

To confirm that the high decodability rate was not an artifact of using all recorded neurons, including those with stark differences in their responses to equally valued safe and risky offers, we used our subpopulation approach. Specifically, we identified putative pure-value cells by taking the set of cells with no significant difference in response to safe and equivalent risky offers (that is, *p* > 0.05). Then, to be even more conservative, we included only half of these cells with the smallest difference between responses to safe and SV-matched risky offers. In the OFC, this constituted 68/157 neurons (43.3% of recorded cells; rOFC: 60/138 neurons, 43.5%; vmPFC: 66/156 neurons, 42.3%; pgACC: 100/255, 39.2%; PCC: 60/151, 39.7%; VS: 50/124, 40.3%). Despite specifically selecting the cells most likely to be abstract value-coding cells, the classifier could readily disambiguate the safe from the SV-matched risky trials (*p* < 0.001; **Figure 4B**). Then, to push even harder against our own conclusions, we conducted the same analysis on an even smaller subset. We included only a quarter of the cells with the smallest difference between responses to safe and SV-matched risky offers. That is, we took the subpopulation that would, by even a very conservative analysis, be most likely to be classified as pure value cells. In the OFC, this constituted 39/157 neurons (24.8% of recorded cells; rOFC: 30/138 neurons, 21.7%; vmPFC: 33/156 neurons, 21.2%; pgACC: 50/255, 19.6%; PCC: 30/151, 19.9%; VS: 25/124, 20.2%). Even so, the classifier could readily disambiguate the safe from the SV-matched risky trials in the remaining five of our structures (*p* < 0.001, in all cases; **Figure 4C**).

### Orthogonal response subspaces for value-matched risky and safe offers

The underlying connectivity of neuron ensembles can effectively constrain activity to be correlated, rendering a low-dimensional dynamics or subspace (Oşan et al., 2007; Gallego et al., 2017; Ebitz and Hayden, 2021). If value is encoded abstractly, the risky and safe offers will necessarily be found within the same *abstract value subspace.* This subspace does not require abstract value neurons. We adapted previously used approaches to characterize the uniqueness of safe and matched-risky subspaces (Elsayed et al., 2016; Yoo & Hayden, 2020). We projected risky offer responses into the safe offer response subspace and computed the explained variance for each (**Figure 5A-F**). We used these projections to quantify the extent to which subspaces were aligned (Aidx; see **Methods**). An alignment index equal to 1.0 would indicate that the variances in risky and safe responses are explained equally well by the principal components that constitute the safe response subspace. That value would suggest common (“aligned”) subspaces and is predicted by the explicit view of value encoding. An alignment index equal to zero would indicate that the risky and safe subspaces are strictly orthogonal and an index between 0 and 1 would indicate a partial orthogonality. Note that even partial orthogonality would indicate distinct organizational principles.

**Figure 5.**
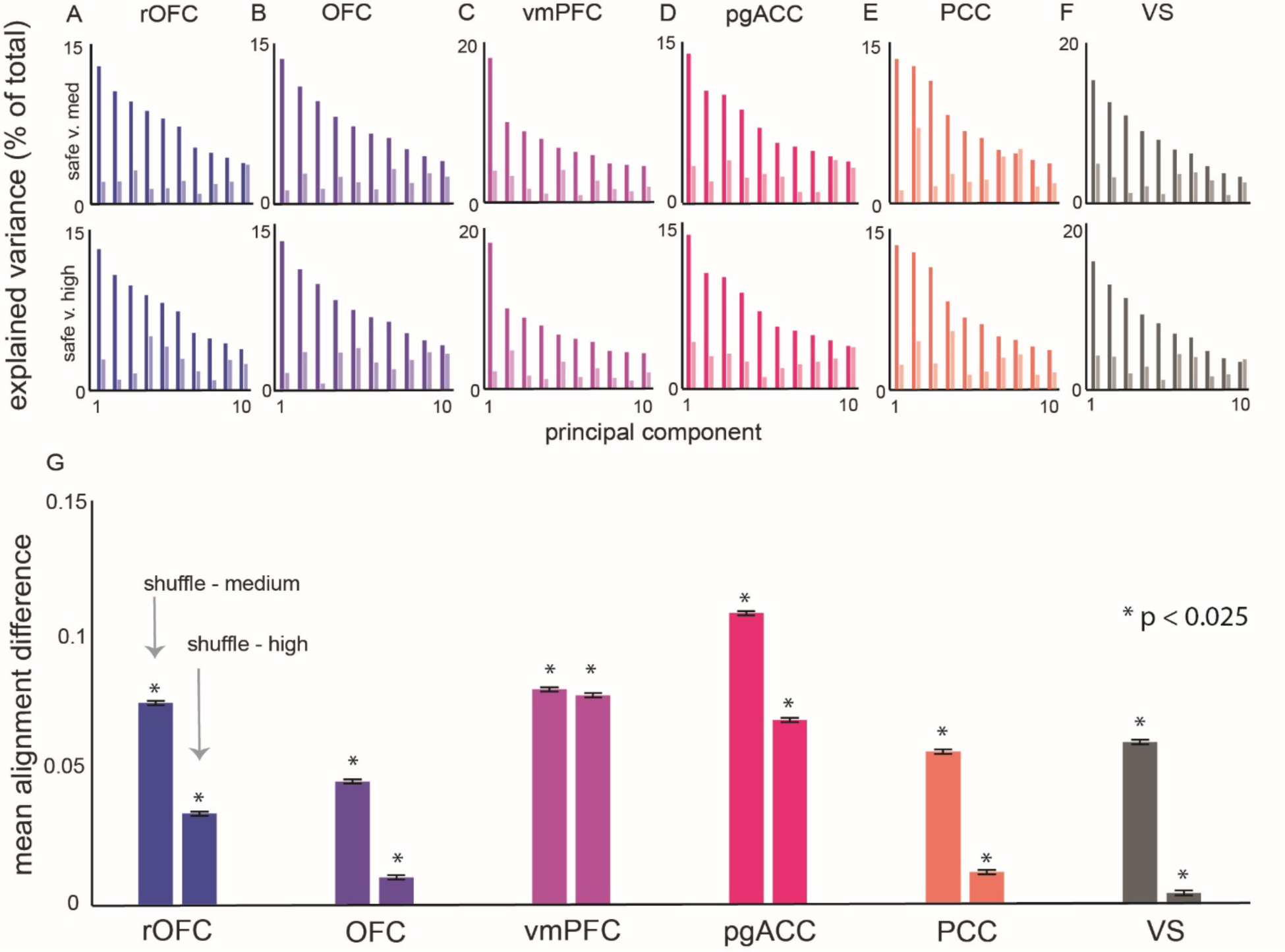
Subspace Alignment between Safe and Equivalent Risky Offers. **(A-F)** For each structure, rOFC, shows the explained variance (as a proportion of total variance). The top panel shows the explained variance in both safe (darker) and medium stakes (lighter) response by projecting both sets of responses into safe response subspace, due to each of the first 10 principal components. The bottom panel is similar, except that it shows explained variance by projecting both safe and high stakes responses into safe response subspace. **(G)** a summary of the difference between the average shuffle alignment index and either the safe-medium (left bar) or the safe-high (right bar) alignment index. Error bars indicate the standard error across computed differences.

The alignment analysis indicated that in OFC, comparisons of safe and medium magnitude risky response subspaces had an alignment index of Aidx = 0.231. Safe and high magnitude subspaces had an Aidx = 0.266. The critical question is whether this value is significantly less than 1.0, which would indicate at least partial orthogonalization. To determine what would constitute a proper threshold for determining partial orthogonalization, we performed a control procedure (see **Methods**) in which we shuffled across time, but not across neurons (Elsayed et al., 2016). Any alignment index at or below this threshold would be considered primarily orthogonal. To do this, we shuffled the data from across all three response matrices (safe, matched-medium, and matched-large) and computed the alignment index between pairs of shuffled sets (see **Methods**). We repeated this process over 1000 randomizations. Then, to test for significance, we computed the 99% confidence interval across iterations; a value outside of this range can therefore be said to be significant at p < 0.005 (given that our t-test is assumed to be one-tailed). We found that the average shuffled alignment index in OFC was Aidx = 0.276. Both the safe-medium and safe-high alignment indexes were below the 99% confidence interval (0.271 - 0.281) and thus both significant at p<0.005 (or p<0.01 with a two-tailed t-test). As a positive control, we repeated the shuffle procedure, but completely randomized across all axes, which provided a relative zero-point for determining strict orthogonality within our data. In OFC, the alignment index across totally shuffled data was Aidx = 0.091 ± 0.009 (99% confidence interval). The safe-medium and safe-high alignment indexes are both quite a bit greater than this noise floor (thus both significant at p<0.005). In other words, response subspaces for safe and equally valued risky offers in OFC are more orthogonal than we would expect by chance, but not perfectly orthogonal, given the inherent statistical properties of our dataset (**Figure 5G**). We found similar results in all structures (below the 99% confidence interval, in all cases; *p* < 0.005).

### Orthogonal risky and safe response subspaces can be transformed to be aligned

Above, we demonstrated that the population subspaces for risky and safe offers are semi-orthogonal (see cartoon in **Figure 6A**). A primary question, then, is to what extent can these subspaces be transformed into a common subspace that would aid in their comparison? Such a common space could provide an underlying, implicit value axis within the population code (Elsayed et al., 2016). We reasoned that if the response subspaces could be transformed such that their hyperplanes were no-longer orthogonal to one another, but aligned, then the axis along which they become aligned is most likely the axis that describes their comparative values (**Figure 6B**). To investigate whether our data here obey these principles, we performed a canonical correlation analysis on the subspace loadings (i.e., the principal component projection weights) for the first three factors from the safe and risky offer response matrices. We also randomly shuffled the subspace loadings 1000 times and performed the same canonical correlation analysis. In OFC, we found that safe and equally valued medium-stakes offer response subspaces could be significantly aligned to correlation of r = 0.53 (*p* = 0.023, bootstrap rank test). Safe and matched-high stakes risky offer response subspaces reached a maximal correlation of Spearman’s correlation of r = 0.61 (*p* < 0.01). We found similar results across the remaining five brain areas in our dataset (*p* < 0.05 in all cases, **Figure 6C-H**). Specifically, we found that in all six brain areas, subspaces were linked.

**Figure 6.**
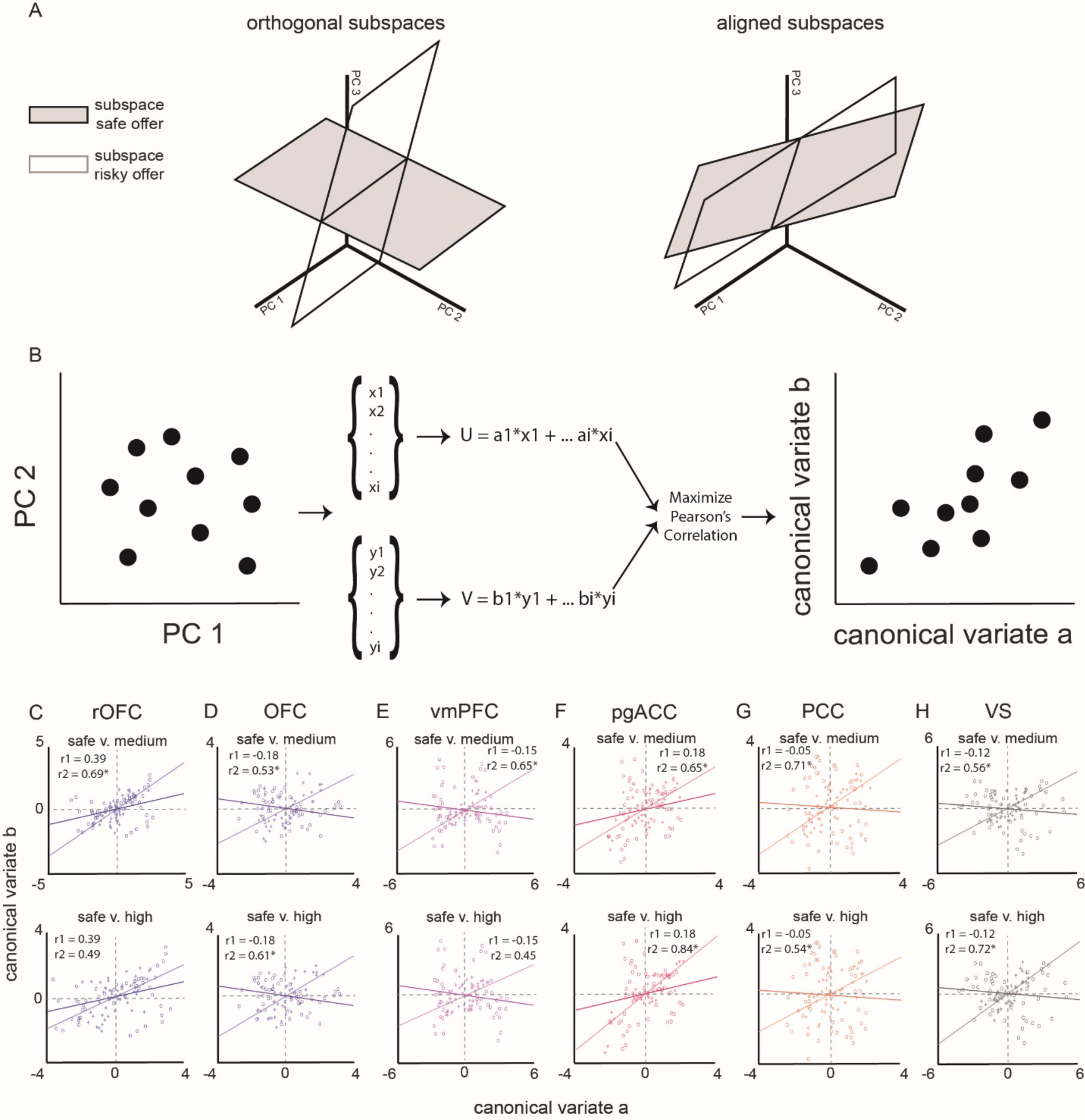
Subspace rotational transformations and canonical correlations. **(A)** A cartoon demonstrating an example of orthogonal (left) and aligned (right) hyperplanes projected onto the first three dimensions. **(B)** A procedural schematic demonstrating the process of linear algebraic rotational transformation, to align subspaces by maximizing the Pearson’s correlation coefficient via canonical correlation. **(C-H)** Scatter plots, for each structure, of projections of medium and safe offer responses (top row) and high and safe offer responses (bottom row) projected into safe offer response subspaces. Plotted are projections onto the first 2 principal component loadings for the original (darker circles) and transformed (lighter pluses) projections. The inlaid coefficients are the Pearson’s correlation coefficients between projections on the first 2 principal component loading for the original (r1) and transformed (r2) responses. The lines denote the best-fit line for the corresponding correlation coefficients (r1: darker line; r2: lighter line).

## DISCUSSION

Our ability to select between qualitatively different options with ease suggests that we can access some kind of value information. The question of how that value is represented in the brain has been a major source of scholarly interest (Tremblay & Schultz, 1999; Elliott et al., 2000; Rangel et al., 2008; Kable & Glimcher, 2009; Levy & Glimcher, 2011; Padoa-Schioppa, 2011; Rushworth et al., 2011; Wunderlich et al., 2012; McNamee et al., 2013). Our findings raise the possibility that abstract value may not be encoded in the activities of single neurons, but instead that value representation may be an emergent process reflecting the coordination activity of distributed sets of expectancy-coding neurons. using the well-established paradigm of risky vs. safe choice, we show that neurons in six core value regions respond to safe offers in a manner that cannot be predicted from their responses to equally valued risky offers. Indeed, the differences in coding of risky and safe offers is highlighted by the fact that they elicit responses in distinct and semi-orthogonal response subspaces. Nonetheless, these subspaces are intrinsically linked, meaning that there is a common subspace into which each can be reliably transformed, which we propose is a shared value subspace that may facilitate comparison and lead to choice. These data support the hypothesis that value coding in the brain is largely implicit and compositional, rather than explicitly coded by single neurons.

We also found that while population responses to the different option types were distinct, they were nonetheless transformable, meaning that their subspaces can be aligned. This arrangement allows the brain, in some sense, to have the best of both worlds. On one hand, the transformability between representations means that they can readily be compared to each other so as to allow for effective economic choice. On the other hand, the difference in coding schemes allows the brain access to information about the properties of the option they are choosing, to know what features to expect, and to adjust their preferences to changing demands (Rigotti et al., 2013; Fusi et al., 2016). For example, if the task suddenly switched so that the probability cues had reversed meanings (and changing rules is a common occurrence in the real world), the decision-maker could rapidly adjust their value response by altering their probability response functions without having to recreate the entire value coding system.

A good deal of evidence, including our own, supports the idea that specialized neurons in the ostensible reward regions of the brain encode value abstractly (e.g., see our claims in Strait et al., 2014, 2015). Our results here are difficult to reconcile with such findings. Specifically, our results indicate that, at least in the domain of risky and safe options, such abstract value neurons are likely to be rare, and indeed, did not appear in measurable quantities in our sample of over 900 neurons. First, we note that, despite the large amount of evidence for abstract value coding, much of it is subject to various forms of confounding, so the case remains open (Maunsell, 2004; Kable & Glimcher, 2009; Wallis & Rich, 2011; O’Doherty, 2014; Stalnaker et al., 2015; Hayden & Niv, 2021; Zhou et al., 2021). We suspect that two main factors differentiate our study from past ones. First, by focusing on qualitatively different types of offers/choices, we bypass the potential confusion caused by mixed selectivity in prefrontal neurons, which can inflate estimates of pure value coding (Fine et al., 2021). Mixed selectivity has long been known, but its implications, and its powerful ability to alter conventional interpretations of physiological analyses, has only recently come to be appreciated (Rigotti et al., 2013; Fusi et al., 2016). Second, risky and safe options are qualitatively different, and evoke different affective responses, whereas most past studies use bundles of goods with similar affective properties (Padoa-Schioppa & Assad, 2006; Lak et al., 2014; Xie & Padoa-Schioppa, 2016; Conen & Padoa-Schioppa, 2019). These results, then, suggest that while neurons in core value regions may correlate with value across some contexts, in the more general case, they fail to satisfy the requirements for abstract value coding. In other words, past studies generally report patterns are that are necessary, but not sufficient, to demonstrate abstract value coding.

While our results do not cohere with past results arguing for pure value coding, they are entirely consistent with a different body of work, one that emphasizes the role of ostensible value regions in encoding expectancy, or the features of the anticipated outcome of a choice (Schoenbaum et al., 2011; Zhou et al., 2021). Expectancy coding has long been associated with the orbitofrontal surface, including rodent homologues of OFC, rOFC, vmPFC, and pgACC (Gardner & Schoenbaum, 2021). It has also been seen as a rival theory to the abstract value theory, and, like that theory, has a great deal of evidence associated with it (Schoenbaum et al., 1998, 2003, 2009, 2011; Zhou et al., 2021). Our results, while they provide clear and direct support for the expectancy side of the theory, offer a potential reconciliation: the expectancy coding found in individual neurons contributes to an implicit value coding. This interpretation, indeed, is supported by our past work on OFC showing that separate dimensions that influence value can use unrelated coding schemes (and by implication, separate subspaces (Blanchard et al., 2015) and that different cues that predict the same reward can elicit unrelated neural responses (Wang & Hayden, 2017). The expectancy theory, more generally, is associated with the idea that the function of OFC is to implement a cognitive map of task space (Wilson et al., 2014; Schuck et al., 2016; Behrens et al., 2018; Schuck & Niv, 2019;). Our results, then, not only endorse the expectancy and cognitive map theories of OFC function, but suggest they may apply to other putative reward regions as well.

A good deal of theorizing holds that the organization of the cortical and striatal reward system is modular. That is, it presumes that value components can be found in hierarchically early regions, and that these are assembled into pure value representations in abstract value areas, typically either OFC, vmPFC, VS, or PCC (Rangel et al., 2008). The fact that we find no abstract value neurons but we do find underlying value coding in all regions suggests that implicit value coding may be the principal way that value is represented in the brain. Moreover, they suggest that value coding is highly distributed (Hunt et al., 2014; O’Doherty, 2014; Hunt & Hayden, 2017; Maisson et al., 2020). Of course, it is possible that abstract value neurons are available in downstream areas, such as DLPFC or dACC. However, neurons in such areas have a clear motor repertoire, which would seem to be inconsistent with the abstract value coding ideas (Tremblay & Schultz, 1999, Pearson et al., 2010, Padoa-Schioppa, 2011). These results, then, raise the intriguing idea that value may be a convenient way to think about brain function, but that its neural implementation may be fundamentally emergent in all areas in which it is found.

## METHODS

### Surgical procedures

All procedures were approved by either the University Committee on Animal Resources at the University of Rochester or the IACUC at the University of Minnesota. Animal procedures were also designed and conducted in compliance with the Public Health Service’s Guide for the Care and Use of Animals. All of the animals were handled according to approved institutional animal care and use committee (IACUC) protocols (#2005- 619 38127A) of the University of Minnesota. The protocol was approved by the Committee on the Ethics of Animal Experiments of the University of Minnesota (NIH permit number: A3456- 01). All surgery was performed under controlled anesthesia, and every effort was made to minimize suffering. Six male rhesus macaques (Macaca mulatta) served as subjects. A small prosthesis head fixation was used. Animals were habituated to laboratory conditions and then trained to perform oculomotor tasks for liquid rewards. We place a Cilux recording chamber (Crist Instruments) over the area of interest (see Behavioral tasks for breakdown). We verified positioning by magnetic resonance imaging with the aid of a Brainsight system (Rogue Research). Animals received appropriate analgesics and antibiotics after all procedures.

Throughout both behavioral and physiological recording sessions, we kept the chamber with regular antibiotic washes and we sealed them with sterile caps.

Data and code availability. Custom code for the following analyses is written in MatLab available through GitHub at https://github.com/dmaisson/DifferentialRiskEncoding. Data used for all reported analyses are available on Dryad (https://doi.org/10.5061/dryad.ttdz08kx9).

### Recording sites

We approached our brain regions through standard recording grids (Crist Instruments) guided by a micromanipulator (NAN Instruments). All recording sites were selected based on the boundaries given in the Paxinos atlas (Paxinos et al., 2008). In all cases we sampled evenly across the regions. Neuronal recordings in OFC were collected from *subjects P and S*; recordings in rOFC were collected from *subjects V and P*; recordings in vmPFC were collected from *subjects B* and *H*; recordings in pgACC were collected from *subject B and V*; recordings from PCC were collected from *subject P and S*; and recording in VS were collected from *subject B and C*. Specifically (see **Figure 1B**):

We defined **rOFC 11** as lying within the coronal planes situated between 34.05 and 42.15 mm rostral to the interaural plane, the horizontal planes situated between 4.5 and 9.5 mm from the brain’s ventral surface, and the sagittal planes between 3 and 14 mm from the medial wall. The coordinates correspond to area 11 in Paxinos et al. (2009).

We defined **OFC 13** as lying within the coronal planes situated between 28.65 and 34.05 mm rostral to the interaural plane, the horizontal planes situated between 3 and 6.5 mm from the brain’s ventral surface, and the sagittal planes between 5 and 14 mm from the medial wall. The coordinates correspond to area 13m in Paxinos et al. (2009).

We defined **vmPFC 14** as lying within the coronal planes situated between 29 and 44 mm rostral to the interaural plane, the horizontal planes situated between 0 and 9 mm from the brain’s ventral surface, and the sagittal planes between 0 and 8 mm from the medial wall. These coordinates correspond to area 14m in Paxinos et al. (2009).

We defined **pgACC 32** as lying with the coronal planes situated between 30.90 and 40.10 mm rostral to the interaural plane, the horizontal planes situated between 7.30 and 15.50 mm from the brain’s dorsal surface, and the sagittal planes between 0 and 4.5 mm from the medial wall (**Figure 1B**). Our recordings were made from central regions within these zones, which correspond to area 32 in Paxinos et al. (2009).

We defined **PCC 29/31** as lying within the coronal planes situated between 2.88 mm caudal and 15.6 mm rostral to the interaural plane, the horizontal planes situated between 16.5 and 22.5 mm from the brain’s dorsal surface, and the sagittal planes between 0 and 6 mm from the medial wall. The coordinates correspond to area 29/31 in Paxinos et al. (2009).

We defined **VS** as lying within the coronal planes situated between 20.66 and 28.02 mm rostral to the interaural plane, the horizontal planes situated between 0 and 8.01 mm from the ventral surface of the striatum, and the sagittal planes between 0 and 8.69 mm from the medial wall. Note that our recording sites were targeted towards the nucleus accumbens core region of the VS (Paxinos et. al, 2009).

We confirmed recording location before each recording session using our Brainsight system with structural magnetic resonance images taken before the experiment. Neuroimaging was performed at the Rochester Center for Brain Imaging on a Siemens 3T MAGNETOM Trio Tim using 0.5 mm voxels. We confirmed recording locations by listening for characteristic sounds of white and gray matter during recording, which in all cases matched the loci indicated by the Brainsight system with an error of ∼1 mm in the horizontal plane and ∼2 mm in the z-direction.

### Electrophysiological techxniques

Either single (FHC) or multi-contact electrodes (V-Probe, Plexon) were lowered using a microdrive (NAN Instruments) until waveforms between one and three neuron(s) were isolated. Individual action potentials were isolated on a Plexon system (Plexon, Dallas, TX) or Ripple Neuro (Salt Lake City, UT). Neurons were selected for study solely on the basis of the quality of isolation; we never preselected based on task-related response properties. All collected neurons for which we managed to obtain at least 300 trials were analyzed; no neurons that surpassed our isolation criteria were excluded from analysis.

### Eye-tracking and reward delivery

Eye position was sampled at 1,000 Hz by an infrared eye-monitoring camera system (SR Research). Stimuli were controlled by a computer running Matlab (Mathworks) with Psychtoolbox and Eyelink Toolbox. Visual stimuli were colored rectangles on a computer monitor placed 57 cm from the animal and centered on its eyes. A standard solenoid valve controlled the duration of juice delivery. Solenoid calibration was performed daily.

### Behavioral tasks

Six monkeys performed in the risky choice task. Both tasks made use of vertical rectangles indicating reward amount and probability. We have shown in a variety of contexts that this method provides reliable communication of abstract concepts such as reward, probability, delay, and rule to monkeys (Blanchard & Hayden, 2015; Sleezer et al., 2016; Mehta et al., 2019; Wang & Hayden, 2019).

### Risky choice task

The task presented two offers on each trial. A rectangle 300 pixels tall and 80 pixels wide represented each offer (11.35° of visual angle tall and 4.08° of visual angle wide; Fig. 2*A*). Two parameters defined gamble offers, *stakes* and *probability*. Each gamble rectangle was divided into two portions, one red and the other either gray, blue, or green. The size of the color portions signified the probability of winning a small (125 μl, gray), medium (165 μl, blue), or large reward (240 μl, green), respectively. We used a uniform distribution between 0 and 100% for probabilities. The size of the red portion indicated the probability of no reward. Offer types were selected at random with a 43.75% probability of blue (medium magnitude) gamble, a 43.75% probability of green (high magnitude) gambles, and a 12.5% probability of gray options (safe offers).

On each trial, one offer appeared on the left side of the screen and the other appeared on the right. We randomized the sides of the first and second offer. Both offers appeared for 400 ms and were followed by a 600-ms blank period. After the offers were presented separately, a central fixation spot appeared, and the monkey fixated on it for 100 ms. Following this, both offers appeared simultaneously and the animal indicated its choice by shifting gaze to its preferred offer and maintaining fixation on it for 200 ms. Failure to maintain gaze for 200 ms did not lead to the end of the trial but instead returned the monkey to a choice state; thus, monkeys were free to change their mind if they did so within 200 ms (although in our observations, they seldom did so). Following a successful 200-ms fixation, the gamble was resolved and the reward was delivered. We defined trials that took > 7 sec as inattentive trials and we did not include them in the analyses (this removed ∼1% of trials). Outcomes that yielded rewards were accompanied by a visual cue: a white circle in the center of the chosen offer. All trials were followed by an 800-ms intertrial interval with a blank screen.

### Estimation of subjective value equivalence

We calculated the indifference point between safe and risky offers. For each subject, independently, we fitted a sigmoidal function to the distribution of choices (safe or risky) across the full range of risky offer probabilities.

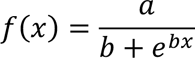

where x = the probability associated with the risky offer, and *f*(x) = the likelihood of choosing the safe offer; a (the maximum value of the curve) and b (the growth rate; steepness) are coefficients of the function, estimated by the fitting procedure for maximizing *R^2^*. Using the fitted sigmoidal function, we then estimated the value of x needed to produce a safe choice likelihood of 0.5; that is, a risky offer choice is equally likely as a safe choice. This is called the *indifference point*, and it allows us to calculate the risky offer value that is considered by the subject to be of equivalent value to that of the safe offer. We performed this analysis separately for medium and high stakes gambles, and separately for each subject.

### Statistical methods

We constructed peristimulus time histograms by aligning spike rasters to the presentation of the first offer and averaging firing rates across multiple trials. We calculated firing rates in 20-ms bins but we analyzed them in longer (500 ms) epochs. Some statistical tests of neuronal activity were only appropriate when applied to single neurons because of variations in response properties across the population. We have used this epoch in all our past research on this and similar tasks (Strait et al., 2014, 2015, 2016; Azab & Hayden, 2017, 2018, 2020). We have found that this epoch provides a good characterization of functional responses and allows for fair comparison across brain regions (Strait et al., 2016; Maisson et al., 2020). We used it here for those reasons and because adherence to a single pre-planned epoch of interest reduces the likelihood of inadvertent p-hacking.

### Evoked response difference analysis

For each neuron, we separated the firing rates in response to offer 1 by whether they corresponded to a safe or risky offer. We calculated the mean firing rates for each neuron across all trials safe and risky trials. We then computed the absolute value of the difference between mean firing rates in response to safe and equivalent risky offers. To determine significance, we performed a 1000-sample bootstrap of randomly selected risky offers, computed the evoked response difference, ranked the evoked response differences in ascending order and asked how many of these 1000 control evoked response differences were less than the true evoked response difference. We next defined subsets of neurons ranging from 1-100% of all recorded neurons in a structure. We performed these same calculations and ordered the differences from least to greatest. For each subset, we took the corresponding number of neurons with the smallest response difference (i.e. the best subset) and compared them to a control population (calculated the same was as described previously). We then performed a Kolmogorov-Smirnov test to compare the average response difference to equal offers across all subset sizes with the response difference to random offers.

### Decoding analysis

We built a pseudo-population of pseudo-trials. First, for each epoch, we isolated firing rate responses to the safe offers and the equivalent risky offers. Then, we collapsed the firing rates for each trial into an average for the 500 ms period. We randomly selected 1000 samples for each neuron, under both risk conditions, resulting in two *n* X 1000 matrices (one for each label level), where n represented the number of neurons recorded from each region. This constituted the pseudo-population of pseudo-trials. To execute the decoder, each matrix was split in half and concatenated with the half from the other label. We used one of these matrices to train a binary support vector machine, the other was used for cross-validation. We used the trained model to predict the binary label (safe or risky) for each pseudo-trial in the cross-validation set. We then compared the predicted label to the known label and an accuracy rate was calculated across predictions. This process was repeated 1000 times for each structure and epoch to get a distribution of accuracy rates. Thus, the standard error of the mean, used in displaying the error bars, represents the standard error over the variance of the cross-validations. Additionally, the exact process was repeated on randomly shuffled data, to confirm that expected prediction accuracy was 50% when randomized.

### Subspace alignment

We followed the procedure described in Yoo & Hayden, 2020. Specifically, for each structure, we separated offers and neural responses by their risk profile (safe and equivalent risky offer of medium and high magnitudes), as described previously. For each neuron, we identified two factors to incorporate into a single condition: time and offer position. Time included the same 500 ms period following the onset of offer 1 and preceding the onset of offer 2. Time was segmented into 20 ms bins. For each 20 ms bin, we computed the mean firing rate across trials on which the offer was positioned on either the left or right of the screen. Thus, we constructed a condition (time X offer position) X neuron matrix of mean firing rates; that is, a 50 X n-neurons matrix. One such matrix was constructed for safe offers, one for medium, and one for high magnitude risky offers of equivalent value. Firing rates in prefrontal areas of macaques tend to be sparse. So, we smoothed these matrices using a gaussian filter, with a sigma equal to one. We then normalized the smoothed matrices, by computing the z-score within each cell, to account for differences in encoding scaling between neurons.

Next, we performed a principal component analysis, using eigenvalue decomposition, on the safe response matrix, providing a transformation matrix into which we projected both the safe response matrix and each of the risky response matrices. We computed the explained variance due to each of the principal components. We performed the same process of dimensionality reduction for each of the risky offers, projecting both the safe response and corresponding risky response data into the resulting principal component spaces (medium magnitude and high magnitude risky response each into their own principal component spaces). To determine if the subspaces were aligned, we computed an alignment index:

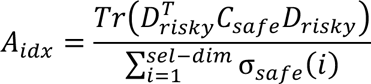

where Tr() is the sum along the diagonal entry, sel-dim = the number of selected principal components (or ten, in the current study), Drisky are the set of top sel-dim eigenvectors, Csafe is the covariance matrix for the safe responses, σsafe(i) is i-th singular value of Csafe. Essentially, the variance explained in safe response by the top ten principal components of the risky responses is normalized against the sum of the variance explained by the top ten principal components of the safe responses. Note that we also performed this calculation using both the top 4 and top 7 principal components. This control did not change the results of the significance tests, and so they are not reported.

To determine the significance of the alignment index we performed a shuffle procedure. We assume, in this procedure, that neural responses adhere to a fixed correlation structure (Elsayed et al., 2016). Thus, we tested whether the safe and risky subspaces were more or less orthogonal, relative to randomly sampling within the space of this fixed correlation structure. We concatenated all safe and risky offer data into a single matrix. We then computed the covariance matrix, and performed an eigenvalue decomposition for the covariance matrix. We then randomly sampled subspaces that were aligned to the fixed correlation structure of the response space, using a method described by Elsayed et al., 2016, as follows:

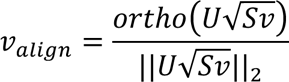

where U and S are the eigenvectors and eigenvalue matrices, respectively, of the computed covariance matrix. A matrix (v) was drawn from a normal distribution with a mean = 0.0 and variance = 1.0. *Orth*() computes the orthonormal basis of the projected matrix. This process essentially maintains the neuronal covariance structure of the original covariance matrix used for the eigenvalue decomposition. We repeated this process across 1000 iterations and computed the alignment index for each, according to the above description. We then calculated the average alignment index and the 95% confidence intervals across the 1000 iterations.

### Multinomial Logistic Regression

To follow-up on the alignment index analysis, we used the projections of safe, medium, and high stakes risky offers into safe subspace. We selected the top 10 results principal components, consistent with the alignment index and constituting no less than 80% of the total explained variance. We then built a multinomial logistic regression model, in which projections onto the first 10 principal components, from of the offer types, were used as simultaneous predictors. The model then set these factors as predictors of a categorical class variable, labeling each offer type. The resulting model was then tested for significance using the standard F-statistic for regressions, against an alpha of 0.05.

## Acknowledgements

We thank Geoffrey Schoenbaum for helpful discussions. We than Marc Mancarella, Caleb Strait and Tommy Blanchard for assistance with data collection, Sarah Heilbronner for help with anatomy, and the rest of the Hayden and Zimmermann labs for valuable discussions. This research was supported by a National Institute on Drug Abuse grant P30 DA048742-01A1 (to BYH and JZ), a National Institute for Biomedical Imaging Grant P41 EB027061 (to BYH and JZ), and a UMN AIRP award (to BYH and JZ).

